# Integrative multiomic approaches reveal ZMAT3 and p21 as conserved hubs in the p53 tumor suppression network

**DOI:** 10.1101/2024.09.17.612743

**Authors:** Anthony M. Boutelle, Aicha R. Mabene, David Yao, Haiqing Xu, Mengxiong Wang, Yuning J. Tang, Steven S. Lopez, Sauradeep Sinha, Janos Demeter, Ran Cheng, Brooks A. Benard, Liz J. Valente, Alexandros P. Drainas, Martin Fischer, Ravindra Majeti, Dmitri A. Petrov, Peter K. Jackson, Fan Yang, Monte M. Winslow, Michael C. Bassik, Laura D. Attardi

## Abstract

*TP53*, the most frequently mutated gene in human cancer, encodes a transcriptional activator that induces myriad downstream target genes. Despite the importance of p53 in tumor suppression, the specific p53 target genes important for tumor suppression remain unclear. Recent studies have identified the p53-inducible gene *Zmat3* as a critical effector of tumor suppression, but many questions remain regarding its p53-dependence, activity across contexts, and mechanism of tumor suppression alone and in cooperation with other p53-inducible genes. To address these questions, we used Tuba-seq^Ultra^ somatic genome editing and tumor barcoding in a mouse lung adenocarcinoma model, combinatorial *in vivo* CRISPR/Cas9 screens, meta-analyses of gene expression and Cancer Dependency Map data, and integrative RNA-sequencing and shotgun proteomic analyses. We established *Zmat3* as a core component of p53-mediated tumor suppression and identified *Cdkn1a* as the most potent cooperating p53-induced gene in tumor suppression. We discovered that ZMAT3/CDKN1A serve as near-universal effectors of p53-mediated tumor suppression that regulate cell division, migration, and extracellular matrix organization. Accordingly, combined *Zmat3*-*Cdkn1a* inactivation dramatically enhanced cell proliferation and migration compared to controls, akin to *p53* inactivation. Together, our findings place *ZMAT3* and *CDKN1A* as hubs of a p53-induced gene program that opposes tumorigenesis across various cellular and genetic contexts.

## Introduction

The *TP53* gene, encoding the p53 protein, is the most frequently mutated gene in human cancer, inactivated in nearly half of tumors (1), underscoring its importance as a tumor suppressor. p53 is a transcription factor which is activated in cells in response to various stress signals and induces transcription of a network of hundreds of target genes encoding proteins involved in diverse biological processes. In human cancer, *TP53* mutations occur predominantly in the DNA binding domain and largely render p53 incapable of activating expression of its target gene network (2). In genetically engineered mouse models (GEMMs), inactivation of both p53 transactivation domains abolishes target gene expression and tumor suppression in a variety of cancer types (3–6). Together, these observations demonstrate the essentiality of transcriptional activation of the network of p53 target genes for tumor suppression. Despite the unequivocal importance of transcriptional activation function for p53-mediated tumor suppression, our understanding of the p53 target genes that directly mediate tumor suppression remains incomplete (7,8). Given the prevalence of *TP53* mutations in cancer and the stark lack of an approved therapy targeting the p53 pathway in the clinic (9,10), a better understanding of the target genes and cellular effects downstream of p53 holds promise for advancing cancer therapy.

In recent years, several p53 target genes have been identified as candidate effectors of p53-mediated tumor suppression *in vivo*, including *Mlh1*, *Abca1*, *Gls2*, *Padi4*, and *Zmat3* (11–15). These genes were identified as functional tumor suppressors using various genetic tools to knock-down, knock-out, and overexpress genes *in vivo*. Importantly, in thinking about the role of p53 target genes in p53-mediated tumor suppression, care must be taken to isolate p53-dependent effects from p53-independent effects. While genetic manipulation of p53 target genes in *p53* wild-type and *p53* null settings has begun to uncover the nuances of p53-dependent versus p53-independent activity, it remains unclear whether abrogating p53-mediated gene induction would phenocopy target protein deletion. Ultimately ablating the response elements (REs) required for p53-dependent induction of specific target genes is critical for unambiguously establishing the contribution of p53-dependent regulation of specific genes for tumor suppression.

Of the recently identified tumor suppressor target genes, *Zmat3*, which encodes an RNA-binding protein that regulates RNA homeostasis (16,17), is notable for having been independently identified as a tumor suppressor in unbiased *in vivo* genetic screens conducted by two different laboratories and for displaying tumor suppressor function in multiple models, including lung adenocarcinoma (LUAD) and hepatocellular carcinoma GEMMs, as well as human colorectal cancer cells (11,15,18). Nonetheless, our understanding of *Zmat3* and other emerging p53 target genes remains preliminary and key questions remain to be addressed. For example, how disruption of p53 regulation of *Zmat3* affects tumor suppression remains unclear. It also remains unclear how much of p53-mediated tumor suppression *Zmat3* carries out and whether *Zmat3* is part of a “core” program of p53 target genes important for p53-mediated tumor suppression conserved across tissue types and species. Beyond these questions regarding *Zmat3* in itself, critical work remains to contextualize *Zmat3* within the broader p53 target gene network. Elucidating which p53 target genes *Zmat3* cooperates with during tumor suppression and which cellular processes and signaling pathways these genes regulate will provide the best strategy for ultimately translating our understanding of the p53 target gene network into therapies that target the p53 pathway for cancer treatment.

In this study, we address these knowledge gaps with an integrative multiomic approach. We first generated *Zmat3* conditional knockout mice for quantitative, multiplexed tumor assays using somatic genome editing and tumor barcoding (Tuba-seq^Ultra^), which revealed a clear p53-dependent role for ZMAT3 in suppressing LUAD, accounting for approximately one third of p53 activity. After establishing ZMAT3 as a major p53 tumor suppressor effector, we next leveraged *in vivo* CRISPR/Cas9 screens to identify p53 target genes cooperating with ZMAT3 in p53-mediated tumor suppression, in which we uncovered *Cdkn1a* as the most potent cooperating tumor suppressor. The importance of *ZMAT3* and *CDKN1A* as hubs in the p53 tumor suppressor network was further supported by our meta-analyses of human gene expression and DepMap functional genetic screening data. Finally, we performed transcriptomic and shotgun proteomic analyses to reveal those aspects of p53-mediated tumor suppression regulated by ZMAT3 and p21. These analyses revealed cell proliferation, migration and extracellular matrix (ECM) signatures induced both upon coordinate *Zmat3;Cdkn1a* inactivation as well as *Trp53* inactivation. Analysis of cellular models showed that, indeed, ZMAT3 and p21 act in a concerted fashion to dampen cellular proliferation and migration in 3D, akin to p53. Collectively, our findings suggest that *Zmat3* and *Cdkn1a* constitute significant effectors of the p53 program, the most significant tumor suppression program in human cancer, thereby revealing fundamental pathways involved in carcinogenesis and paving the way toward the identification of new targets for cancer therapy.

## Results

### Tuba-seq^Ultra^ provides a quantitative measurement of p53-dependent tumor suppression by *Zmat3 in vivo*

A growing body of evidence supports the importance of *Zmat3* in p53-mediated tumor suppression (11,15,18). However, the extent of the contribution of ZMAT3 to tumor suppression relative to p53 itself and which aspects of tumor suppression ZMAT3 modulates, such as tumor initiation and/or growth, remain to be established. Furthermore, the precise p53-dependent contribution of *Zmat3*, as modeled by disrupting the *Zmat3* p53 response element (RE), is unknown. To address these questions, we employed somatic genome editing, tumor barcoding, and high through-put barcode sequencing (Tuba-seq^Ultra^) in an autochthonous KRAS^G12D^-driven, mouse LUAD model where *Zmat3* has tumor suppressor function (15,19–21). We generated a pool of barcoded Lenti-U6^BC^-sgRNA/Cre vectors that included vectors expressing sgRNAs targeting *Trp53*, *Zmat3*, or the p53 response element in *Zmat3* Intron 1 (*Zmat3* RE), with 3 different sgRNAs per target, as well as vectors expressing inert control sgRNAs (Fig. 1A). The *Zmat3* RE sgRNAs targeted a p53 RE which we showed previously to be required for *Zmat3* induction by p53 in mouse embryonic fibroblasts (MEFs; Bieging-Rolett et al. 2020). Our sgRNA pool allowed us to quantify and compare the effects of *Trp53* and *Zmat3* inactivation (Fig. 1Bi) as well as to disentangle p53-dependent (Fig. 1Bii) and p53-independent (Fig. 1Biii) effects of *Zmat3* by comparing the *Zmat3* sgRNAs with those targeting the p53 RE in *Zmat3*.

**Figure 1.**
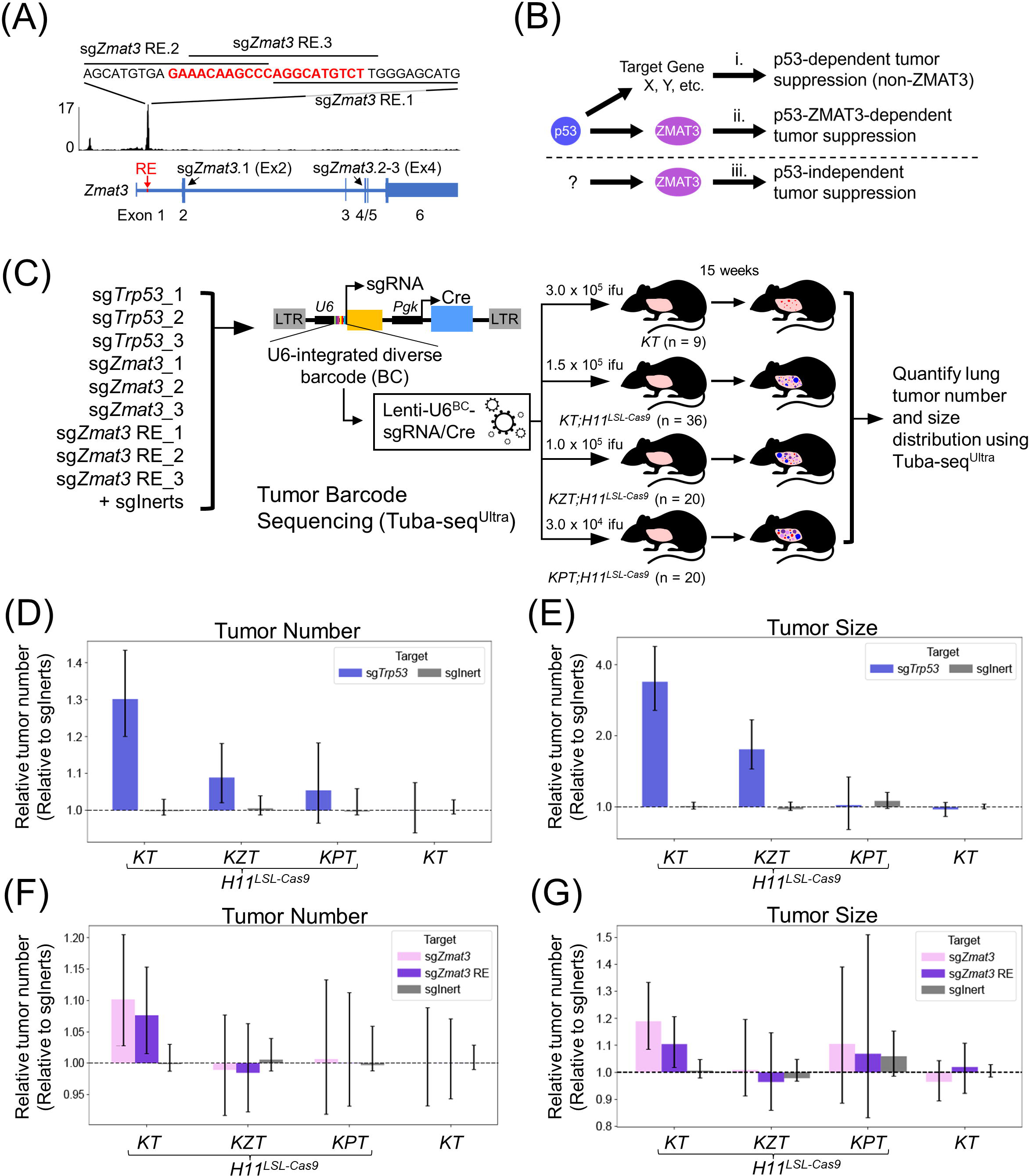
Genetic epistasis mapping reveals the effects of the p53-*Zmat3* LUAD suppression axis *in vivo*. **(A)** Murine *Zmat3* p53 response element (RE) with sgRNA for both sg*Zmat3* RE and gene-targeting sg*Zmat3*. **(B)** Schematic of p53-mediated and ZMAT3-mediated tumor suppression, depicting ZMAT3-independent effects of p53 (i), p53-ZMAT3 effects (ii), and p53-independent effects of ZMAT3 (iii). **(C)** Design of tumor barcoding sequencing (Tuba-seq^Ultra^) experiment. Each gene was targeted with 3 sgRNA, and inert, negative control (sgInert) were included. The lentiviral vector delivers a U6-barcoded sgRNA (BC-sgRNA) and *Pgk*-Cre. Indicated viral titers (infectious units; ifu) of the lentiviral barcoded sgRNA library were delivered to the indicated genotypes and tumorigenesis proceeded for 15 weeks before Tuba-seq analysis was carried out. **(D)** Relative tumor number for sg*Trp53.* **(E)** Log-normal mean tumor size using adaptive cutoff for sg*Trp53*. **(F)** Relative tumor number for sg*Zmat3* and sg*Zmat3* RE. Error bars indicate the 95% confidence interval. sgInert data graphed are aggregate of 3 arbitrarily chosen sgInerts. **(G)** Log-normal mean tumor size using adaptive cutoff for sg*Zmat3* and sg*Zmat3* RE.

To comprehensively investigate the p53-*Zmat3* axis, we initiated lung tumors with this Lenti-U6^BC^-sgRNA/Cre pool in *Kras^LSL-G12D/+^;Rosa26^LSL-tdTomato^* (*KT*), *KT;H11^LSL-Cas9/LSL-Cas9^* (*KT;H11^LSL-Cas9^*), *K;Zmat3^fl/fl^;T;H11^LSL-Cas9^* (*KZT;H11^LSL-Cas9^*), and *K;Trp53^fl/fl^;T;H11^LSL-Cas9^* (*KPT;H11^LSL-Cas9^*) mice (Fig. 1C). To establish the *KZT;H11^LSL-Cas9^* cohorts, we generated *Zmat3* conditional knockout mice, in which exons 4 and 5 of *Zmat3* are flanked by loxP sites, and in which we validated Cre-mediated recombination and ZMAT3 inactivation (Supplementary Fig. 1). The *KT;H11^LSL-Cas9^* mice provided a baseline for phenotypes caused by different sgRNAs while *KZT;H11^LSL-Cas9^* mice allowed us to quantify the effects of *Trp53* deletion in the absence of its key effector *Zmat3.* The *KPT;H11^LSL-Cas9^* mice allowed us to uncover any p53-independent effects of *Zmat3* ablation. Finally, *KT* mouse tumors provided key controls for assessing sgRNA library representation and Cas9-dependency of phenotypes (see Methods).

We next used the Tuba-seq^Ultra^ analysis pipeline to identify the effects of sgRNAs on tumorigenesis within and across the genotypes. To this end, DNA was extracted from tumor-bearing lungs 15 weeks after tumor initiation, and the BC-sgRNA region was PCR amplified and sequenced. Analysis of the number of barcodes associated with each sgRNA and the number of reads of each BC-sgRNA enabled the calculation of the effect of each sgRNA on tumor number and tumor size, respectively. To validate the system, we first examined the effects of sg*Trp53* on tumors. We found that sg*Trp53* increased tumor number relative to control sgRNAs in *KT;H11^LSL-Cas9^* mice, indicating a role for p53 in LUAD initiation (Fig. 1D). Next, to examine tumor size, we used adaptive sampling (see Methods) when calculating the log-normal mean tumor size, based on previous data for the phenotype of *Trp53* inactivation in this model (22). We found that in *KT;H11^LSL-Cas9^* mice, sg*Trp53* increased log-normal mean tumor size, as well as the size of the largest tumors, relative to control sgRNAs (Fig. 1E, Supplementary Fig. 2A). Collectively, these observations align with previous findings, both providing validation of the system and supporting the role of p53 in suppressing LUAD initiation and growth (20,23,24). As expected, all tumor promoting effects of sg*Trp53* were ablated in *KPT;H11^LSL-Cas9^*mice (Fig. 1D-E, Supplementary Fig. 2A).

We next quantified the contribution of *Zmat3* to tumor suppression. To this end, we first assessed the effect of sg*Trp53* in *KZT;H11^LSL-Cas9^*mice. Interestingly, the effects of sg*Trp53* expression on tumor number and tumor size were significantly attenuated in *KZT;H11^LSL-Cas9^* mice relative to *KT;H11^LSL-Cas9^* mice (Fig. 1D-E, Supplementary Fig. 2A), bolstering the idea that *Zmat3* is a critical LUAD suppressor downstream of p53 signaling. In further support of this idea, both sg*Zmat3* and sg*Zmat3* RE significantly enhanced tumor number and size in *KT;H11^LSL-Cas9^* mice, but did not do so in either *KZT;H11^LSL-Cas9^*or *KPT;H11^LSL-Cas9^* mice, where *Zmat3* should not be expressed (Fig. 1F-G, Supplementary Fig. 2B), supporting a role for *Zmat3* in p53-dependent suppression of both LUAD initiation and growth. *Zmat3* and *Zmat3* RE deletion in *KT;H11^LSL-Cas9^* mice produced indistinguishable effects, further suggesting that the tumor suppression effects of *Zmat3* are highly p53-dependent in this model (Fig. 1F-G). Collectively, these data suggest that *Zmat3* is a strictly p53-dependent suppressor of KRAS-driven LUAD, representing, conservatively, about one third of p53-mediated tumor suppression in this context, an estimate derived by comparing the effect of sg*Trp53* and sg*Zmat3*/sg*Zmat3* RE on tumor size and number in *KT;H11^LSL-Cas9^* mice (Fig. 1D-G).

### Mapping cooperation in the p53 tumor suppressor network *in vivo*

Our Tuba-seq^Ultra^ experiment underscores the importance of p53-dependent *Zmat3* expression in p53-mediated tumor suppression. These data also support previous findings that *Zmat3* inactivation does not match the full effect of *Trp53* inactivation deletion on promoting tumorigenesis, suggesting that *Zmat3* cannot act alone downstream of p53. Thus, to map critical tumor suppression effectors downstream of p53 that cooperate with *Zmat3* in tumor suppression, we designed an *in vivo* functional genetic screen to assess tumor suppressor activity of direct p53-regulated genes in both a wild-type background as well as a *Zmat3*-deficient background. We selected *E1A;Hras^G12V^*-expressing primary MEFs as a tumor model because it relies on primary cells with an intact p19^ARF^-p53 tumor suppression axis and because of the dramatic suppressive effect of p53 on tumor growth in this model (25,26). We integrated RNA-sequencing (seq) data from *E1A;Hras*^G12V^;*H11^Cas9^ Trp53* wild-type and *Trp53* null MEFs cultured at physiological oxygen (5%) with MEF p53 ChIP-seq data to identify 272 direct, activated p53 target genes (Supplementary Table 1) as candidate tumor suppressors in these cells (Fig. 2A) (27). We next generated an sgRNA library with 10 sgRNAs targeting each gene and 1000 negative control sgRNAs, which we lentivirally transduced into *E1A;Hras*^G12V^;*H11^Cas9^* MEFs (WT screen; Fig. 2B, Ci). We validated both Cas9 cutting efficiency and p53 activity in these cells (Supplementary Fig. 3A,B), verified representation of sgRNAs at the time of injection (T0) (Supplementary Fig. 3C), and grew tumors for 3 weeks after grafting transduced cells subcutaneously into immunocompromised mice. To identify p53 target genes that cooperate with *Zmat3* in tumor suppression, we repeated the screen in parallel using *Zmat3*-KO cells (*Zmat3*-KO screen, Fig. 2Cii), along with a third screen with *p53*-KO cells, which allowed us to confirm the p53-dependence of any hits scoring as tumor suppressive (*p53*-KO screen, Fig. 2Ciii). Tumors from *p53*-KO screen cells were the largest, followed by *Zmat3*-KO screen tumors, and then the WT screen tumors (Fig. 2D), again confirming ZMAT3 as a tumor suppressor but not as potent as p53.

**Figure 2.**
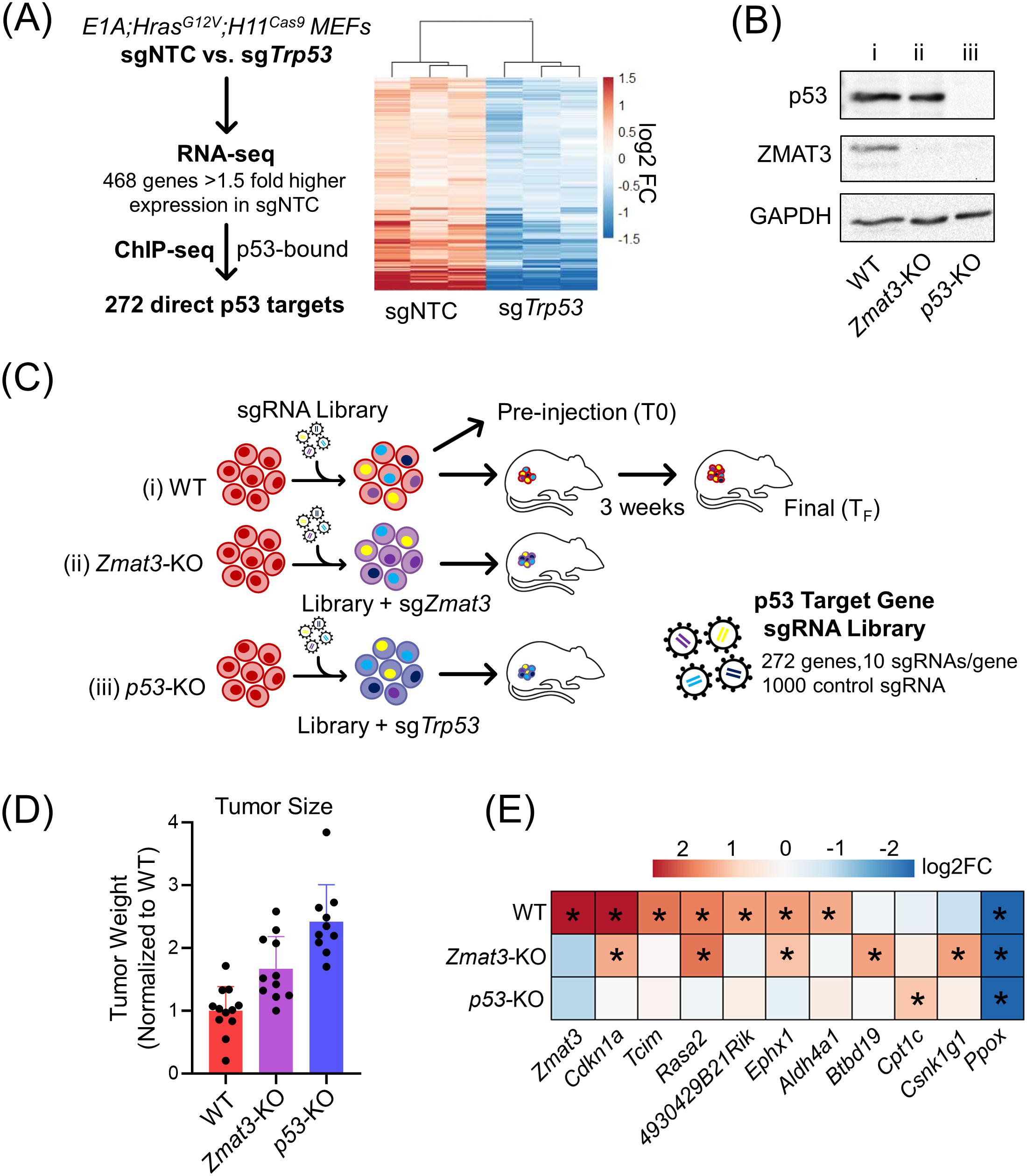
An *in vivo* CRISPR/Cas9 screen uncovers effectors of p53-mediated tumor suppression. **(A)** (Left) Design for sgRNA library targeting direct, upregulated p53 target genes and (Right) heat map showing expression of 272 direct p53 target genes in non-targeting control sgRNA (sgNTC) versus sg*p53 E1A;Hras^G12V^;H11^Cas9^* MEFs. **(B)** Immunoblot of p53, ZMAT3, and loading control (GAPDH) for transduced, selected, pre-injection cell populations for wild-type (WT, i), *Zmat3-*KO (ii), and *p53-*KO (iii) screens. **(C)** p53 target gene sgRNA library was lentivirally delivered to *E1A;Hras*^G12V^;*H11^Cas9^* MEFs, which were grafted subcutaneously in immunocompromised mice to form tumors. Enrichment of sgRNAs was determined by comparing sgRNA representation in final tumors (T_F_) to pre-injection (T0) cells. *Zmat3*-KO (ii) *-* and *p53-* KO (iii) screens were performed in the same way as the WT screen (i) but the sgRNA library was co-transduced with sg*Zmat3* or sg*p53*, respectively. **(D)** Final tumor weight (n = 8-10) of *in vivo* screen samples normalized to wild-type (WT) average. **(E)** Heat map of log2 fold change screen phenotype of genes across all 3 screens from MEMcrispR analysis. *MEMcrispR FDR < 0.05.

We next identified tumor suppressor genes by quantitatively measuring sgRNAs enriched in tumors (T_F_) relative to T0 (Fig. 2E). To this end, we used two algorithms – casTLE (28) and MEMcrispR (29) – to most accurately identify putative tumor suppressors (Fig. 2E, Supplementary Fig. 4A,B, Supplementary Table 2). Strikingly, in this comprehensive pool of p53 target genes in MEFs, *Zmat3* was once again the top hit in the WT screen, underscoring its centrality to p53-mediated tumor suppression (Fig. 2E). The classical p53 target gene *Cdkn1a*, which encodes the cyclin-dependent kinase inhibitor p21 (30), was identified as the second most potent hit in the WT screen, suggesting that *Cdkn1a* is also a critical effector of p53-mediated tumor suppression (Fig. 2E). The *Zmat3*-KO screen revealed 4 enriched hits: *Rasa2*, *Csnk1g1*, *Btbd19*, and *Cdkn1a* (Fig. 2E, Supplementary Fig. 4C,D, Supplementary Table 3), indicating that inactivation of any of these genes can cooperate with *Zmat3* deficiency to promote tumorigenesis. Importantly, analysis of the *p53*-KO screen showed minimal p53-independent effects. Only sg*Cpt1c* was mildly enriched in the *p53*-KO screen according to both casTLE and MEMcrispR, suggesting that all other tumor suppressive genes identified from the WT and *Zmat3*-KO screens have largely p53-dependent effects (Fig. 2E, Supplementary Fig. 4E, F, Supplementary Table 4). Together, these 3 complementary *in vivo* screens underscore the importance of *Zmat3* specifically in p53-driven tumor suppression and identify genetic cooperators with *Zmat3* in tumor suppression.

### *ZMAT3* is a core effector of human p53-mediated growth suppression

Having mapped key target genes that cooperate with *Zmat3* in p53-mediated tumor suppression, we next aimed to assess whether *Zmat3* and other hits from the WT and *Zmat3*-KO screens are members of the “core” program of p53 target genes, genes that are widely bound and upregulated transcriptionally by p53 across tissue, genetic, and evolutionary contexts. To this end, we performed a meta-analysis of 57 human datasets testing p53-dependent gene expression in various cell types treated with diverse stimuli (Fig. 3A; Fischer et al. 2022). Notably, *ZMAT3* and *CDKN1A* were significantly p53-activated in all 57 datasets from a range of human cell types, including colorectal, breast and bone cancer cell lines as well as primary cells. In comparison, analysis of *Zmat3*-cooperators *BTBD19* and *CSNK1G1* revealed broad but not universal p53-dependent gene expression, while *ALDH4A1*, *EPHX1*, *RASA2*, and *TCIM* were more sporadically p53-activated (Fig. 3A). Analysis of 28 human p53 ChIP-seq data sets showed that *ZMAT3* and *CDKN1A* are nearly universally p53-bound in varied human cell types, whereas other screen hits are only occasionally p53-bound (Fig. 3B). In 15 mouse gene expression data sets (31), p53-dependent expression was widespread for *Zmat3*, *Cdkn1a*, and *Btbd19* (Fig. 3C), while meta-analysis of mouse p53 ChIP-seq data sets identified p53-binding peaks for *Zmat3*, *Cdkn1a*, and *Csnk1g1* in all studies (Fig. 3D). Thus, we characterize *Zmat3* and *Cdkn1a* as part of the core p53 program, while other hits from our screen likely play more limited, context-dependent roles in p53-mediated tumor suppression. *ZMAT3* and *CDKN1A* also show broader p53-dependence across contexts than other classical p53 target genes [e.g. *BAX, PMAIP1* (*NOXA*), and *BBC3* (*PUMA*)] (Fig. 3A,B). Collectively, these data position *ZMAT3* and *CDKN1A* as two of the most conserved p53 target genes across cellular and evolutionary contexts with a role in tumor suppression.

**Figure 3.**
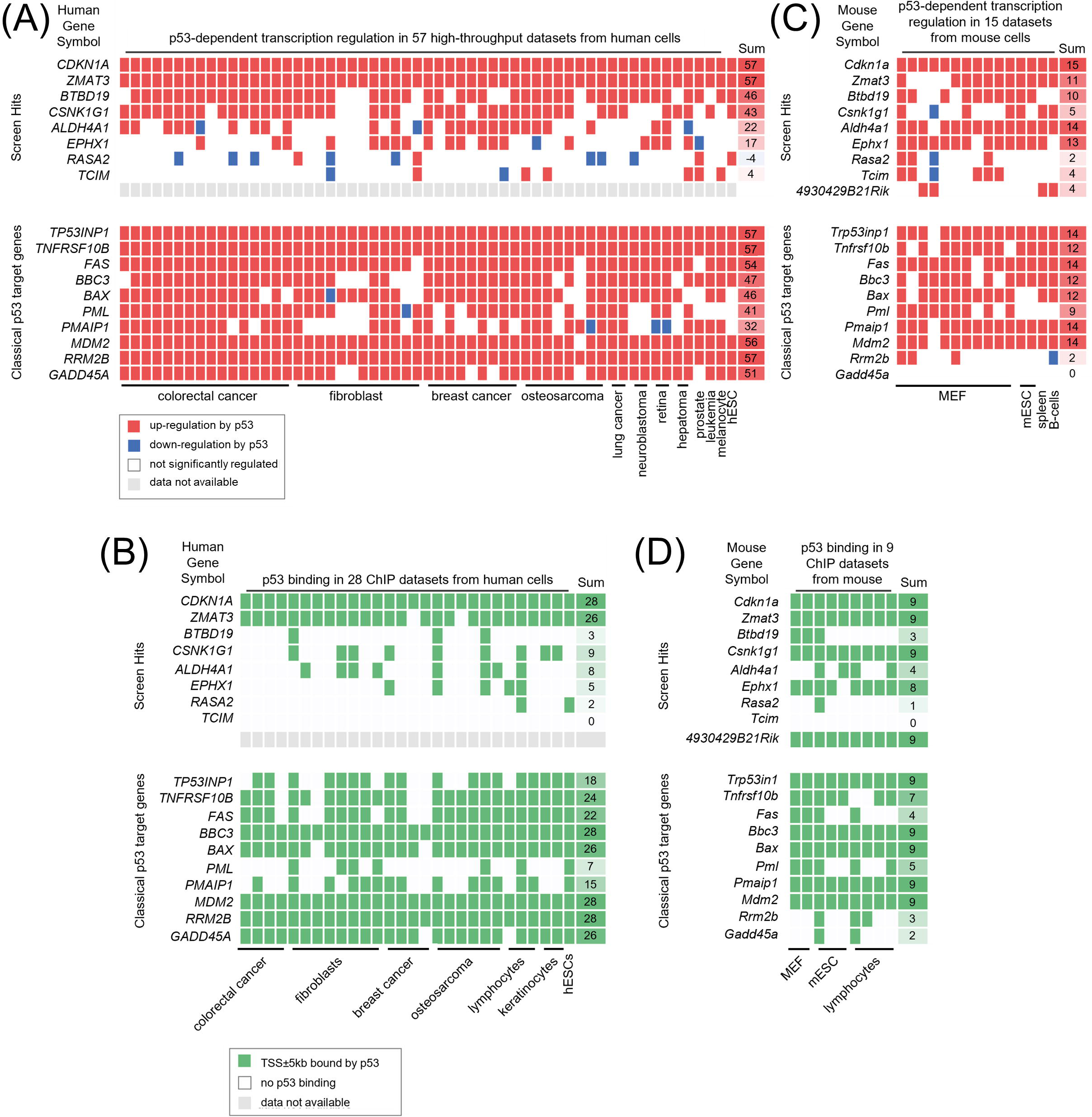
Meta-analysis of p53-dependent expression data from mouse and human cells. **(A)** p53-dependent expression from 57 human datasets. **(B)** Presence of a p53 binding event within 5kb of the transcription start site (TSS) in 26 ChIP-seq datasets from human cells. **(C)** p53-dependent expression from 15 mouse datasets. **(D)** Presence of a p53 binding event within 5kb of the transcription start site (TSS) in 9 datasets from mouse cells. *4930429B21Rik* has no clear human ortholog.

Next, to interrogate the functional importance of p53 target genes in growth suppression of human cells, we performed a pan-cancer analysis of Cancer Dependency Map (DepMap) data, in which genome-wide CRISPR/Cas9-mediated knockout screens were used to assess the effects of gene knockout on cellular fitness in >2,000 human cell lines. In these screens, genes receive a positive gene effect score when their knockout enhances cellular fitness (i.e. increases proliferation, reduces cell death), while a negative gene effect score reflects a reduced cellular fitness phenotype when that gene is knocked out. Using mutation and copy-number status, we stratified cell lines as either *TP53* wild-type or deficient and then calculated the difference in mean gene effect score between *TP53* wild-type and deficient lines for all genes (ΔEffect; Fig. 4A, Supplementary Table 5). Confirming the validity of the analysis, the genes with the greatest ΔEffect between *TP53* wild-type and *TP53*-deficient cell lines were *TP53* and the negative regulator of p53, *MDM2*, as expected (Fig. 4B, Supplementary Table 5). Other well-established upstream activators of the p53 pathway, such as *TP53BP1* and *USP28*, also ranked at the top of the list of genes with the greatest ΔEffect (Fig. 4B, Supplementary Table 5). Analysis of 343 p53 target genes defined by meta-analysis of 17 p53-dependent gene expression studies in human samples (32) revealed *ZMAT3*, along with *CDKN1A*, as the top p53 target genes with positive gene effect scores when ranked by ΔEffect (Fig. 4C, Supplementary Table 6). This combination of positive gene effect score and large ΔEffect indicates a strong tumor suppressor function downstream of p53. In contrast, many classical p53 target genes failed to achieve significant ΔEffect in this analysis (Fig. 4D, Supplementary Table 6). These DepMap data position *ZMAT3* and *CDKN1A* as central components of p53-mediated growth suppression across human cells derived from a wide range of tissue types. Consistent with these genetic findings, a pan-cancer differential gene expression analysis using DepMap data confirmed that *ZMAT3* and *CDKN1A* are among the top genes displaying significantly higher expression in *TP53* wild-type cells than in *TP53-*deficient cells (Fig. 4E, F, Supplementary Table 7). These expression and phenotypic data from DepMap, together with our meta-analysis of p53-dependent gene expression across mouse and human cell lines, highlight the importance of both *ZMAT3* and *CDKN1A* in the p53 tumor suppressor pathway in both mice and humans and support the notion that the most central aspects of p53-mediated tumor suppression are evolutionarily conserved between mouse and human (7,33).

**Figure 4.**
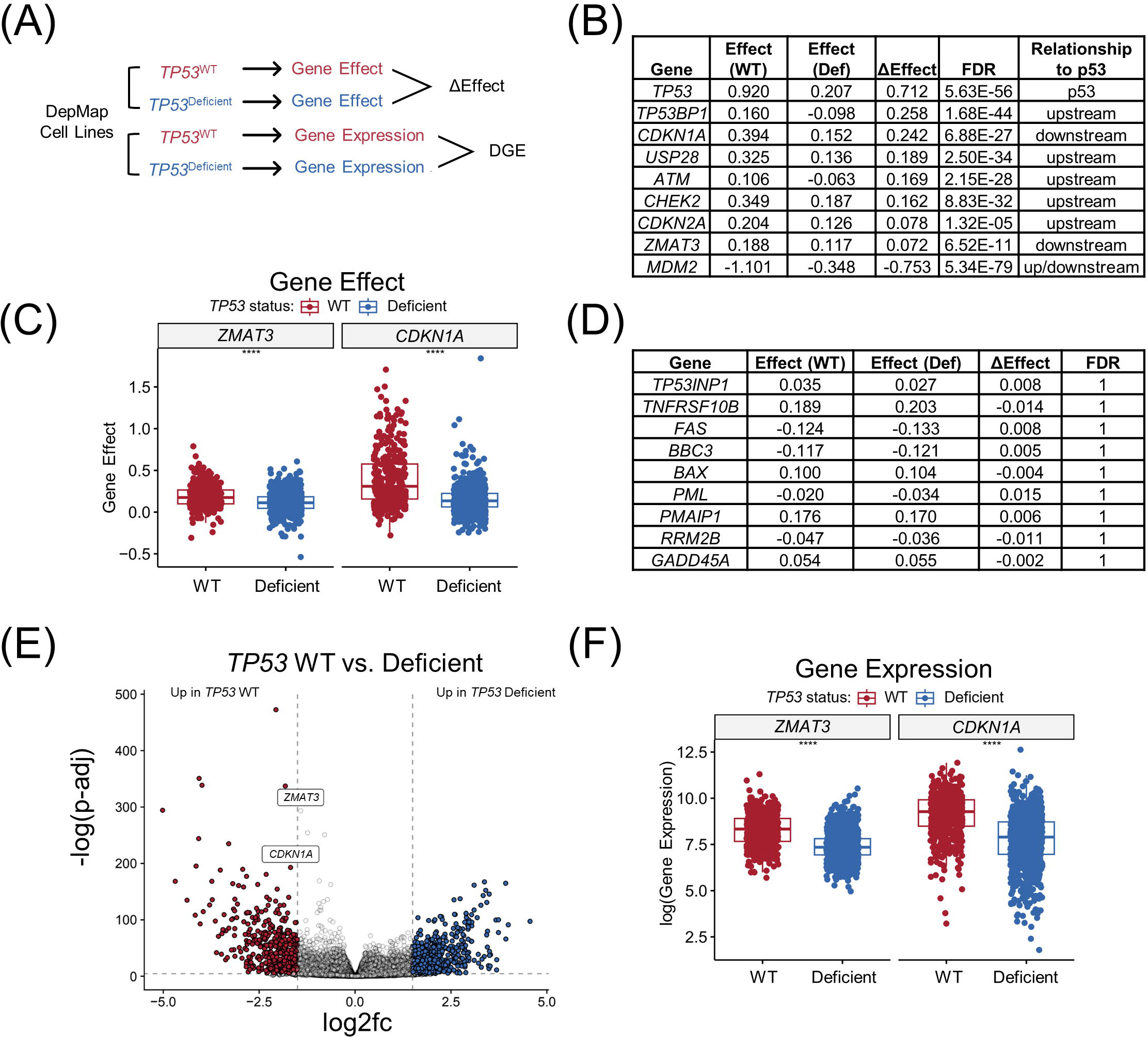
*ZMAT3* and *CDKN1A* are essential and evolutionarily conserved tumor suppressors in the p53 target gene network. **(A)** Schematic of DepMap analyses for *TP53* wild-type (WT) compared to *TP53-*deficient cell lines for both gene effect score (ΔEffect) and differential gene expression (DGE). **(C)** Table of mean gene effect score in *TP53* WT versus *TP53-*deficient (Def) cells for key components upstream and downstream of p53 in the p53 pathway. ΔEffect equals the mean gene effect score in *TP53* WT minus the mean gene effect in *TP53*-deficient cells. False discovery rate (FDR) was calculated for the ΔEffect. **(D)** Plot of gene effect score for *ZMAT3* and *CDKN1A* in *TP53* WT and *TP53-*deficient cell lines in DepMap. ****p < 0.0001. **(E)** Table of mean gene effect score in *TP53* WT versus *TP53-*deficient cells for classical p53 target genes. ΔEffect equals the mean gene effect score in *TP53* WT minus the mean gene effect in *TP53*-deficient cells. False discovery rate (FDR) was calculated for the ΔEffect. **(F)** Volcano plot of differentially expressed genes in *TP53* WT compared to *TP53*-deficient DepMap pan-cancer cell lines. log2fc significance shown at 1.5. **(G)** Plot of mean gene expression for *ZMAT3* and *CDKN1A* in *TP53* WT and *TP53-*deficient cell lines in DepMap. ****p < 0.0001. A two-tailed Student’s t-test was used in (D) and (G).

### *Zmat3* and *Cdkn1a* deficiency drive cancer-associated gene expression programs

Having identified *Zmat3* and *Cdkn1a* as p53 target genes with conserved tumor suppressor function, we next sought to understand the molecular consequences of their deletion by characterizing gene expression profiles. We conducted RNA-seq and shotgun proteomics on control, *Zmat3-*KO, *Cdkn1a-*KO, *Zmat3* + *Cdkn1a* double-KO (DKO), and *p53-*KO *E1A;Hras*^G12V^;*H11^Cas9^* MEFs to elucidate gene expression programs involved in tumor phenotypes (Fig. 5A,B, Supplementary Table 8-9). Differential gene expression analysis of *Zmat3*-KO cells and control cells yielded 3742 differentially expressed genes (padj < 0.05, fold change > 1.5), while *Cdkn1a*-KO cells did not show dramatic changes in gene expression compared to controls with only 43 significantly differentially expressed genes (Supplemental Table. 8). The 4305 differentially expressed genes defined by comparing DKO cells and control cells had striking overlap with *Zmat3*-KO cells (Supplemental Fig. 5A), although pairwise comparison of *Zmat3*-KO cells and DKO cells still revealed 384 significantly differentially expressed genes (Supplemental Table 8). Given the canonical role of p21 in cell cycle regulation, we hypothesized that DKO cells would display enhanced expression of proliferative signatures relative to *Zmat3*-KO cells. Indeed, functional annotation of the 136 upregulated genes in DKO cells versus *Zmat3*-KO cells revealed that “cell division” was the most significant category describing these genes, and was accompanied by other cell cycle categories, consistent with the well-documented role of p21 regulating the cell cycle (Supplemental Fig. 5B).

**Figure 5.**
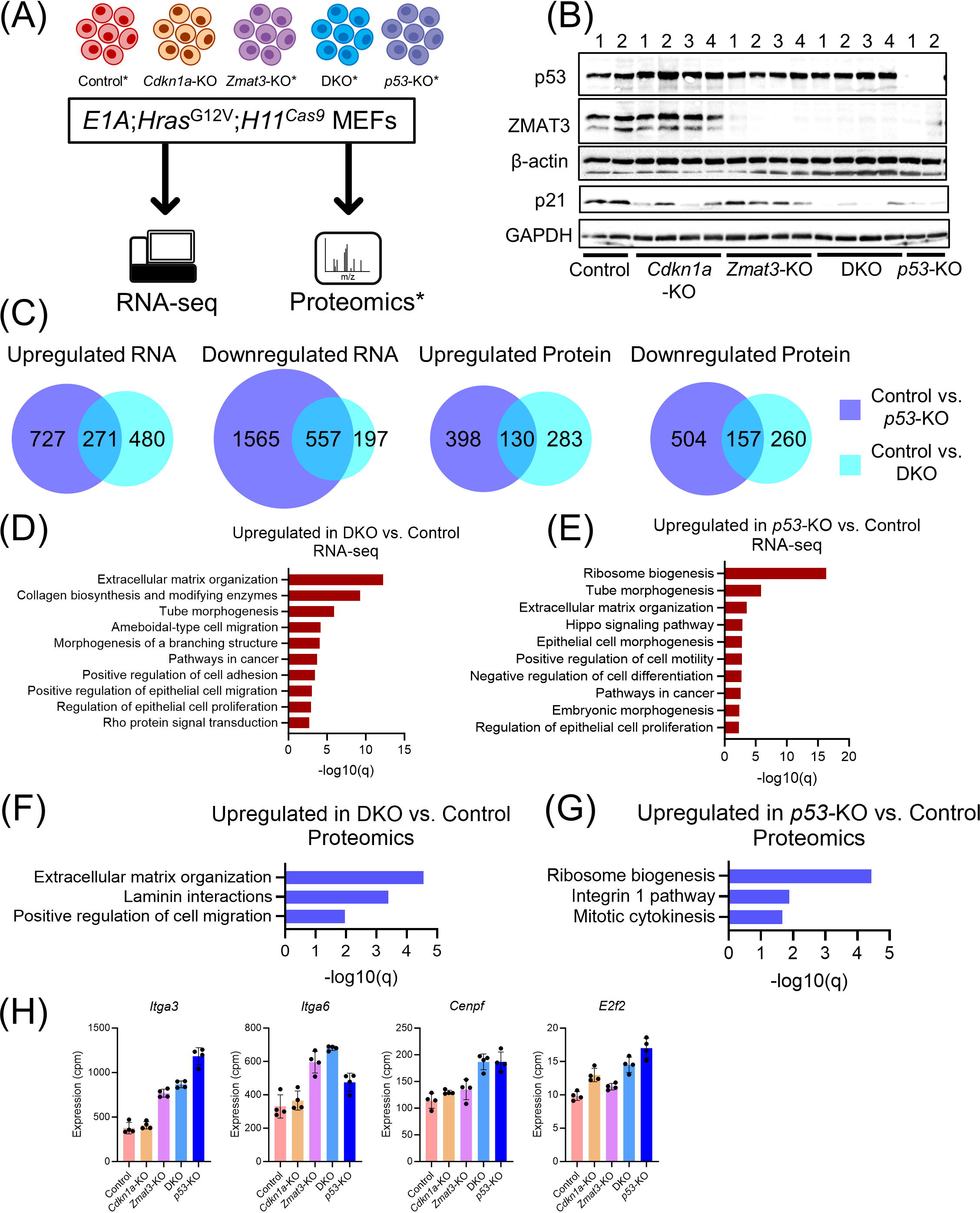
Molecular analysis of genetic cooperation in the p53 tumor suppression pathway. **(A)** Schematic of cell lines used for RNA-sequencing and shotgun proteomic analyses. All genotypes were assessed by RNA-seq. Shotgun proteomics analyses were also conducted for genotypes indicated with *. **(B)** Western blot of samples of cells used for RNA-seq experiment probed for p53, ZMAT3, and p21 as well as loading controls (ACTB and GAPDH). Numbers denote the replicates for each genotype as indicated in the Methods. **(C)** Overlapping significantly differentially expressed genes for pairwise Control vs. *p53*-KO and Control vs. DKO comparisons. Significance was called as padj < 0.05 and fold-change > 1.5 for RNA and padj < 0.05 for protein. **(D,E)** Metascape analysis showing functional annotation for significantly upregulated transcripts in pairwise comparisons of RNA-seq data. Significance was called as padj < 0.05 and fold-change > 1.5. **(F,G)** Metascape analysis showing functional annotation for significantly upregulated proteins in pairwise comparisons of shotgun proteomics data. Significance was called as padj < 0.05. **(H)** Expression for genes associated with cell migration (*Itga3, Itga6*) and cell division-associated genes of interest (*Cenpf*, *E2f2*) identified from RNA-seq (cpm).

Because ZMAT3 and p21 appear to represent a sizable portion of p53-mediated tumor suppression, we reasoned that the molecular events and gene expression changes shared by DKO and *p53*-KO cells can highlight processes most salient to tumor suppression. To this end, pairwise analysis of RNA-seq data indicated notable overlap between genes upregulated and downregulated in both DKO and *p53*-KO cells compared to control cells (Fig. 5C). To highlight oncogenic pathways that might serve as therapeutic targets, we focused our analysis on genes upregulated in DKO and *p53*-KO relative to controls. Functional annotation of such enriched transcripts revealed top categories of gene expression programs involved in cell adhesion, tissue morphogenesis, extracellular matrix (ECM) organization, and other signaling pathways relevant to cell motility and invasion (Fig. 5D,E, Supplementary Table 10). We also found that genes associated with epithelial proliferation were upregulated in both DKO and p53-KO cells, supported by our observation that both DKO and *p53*-KO cells show enhanced proliferation relative to control cells *in vitro* (Supplemental Fig. 5C). Interestingly, functional annotation of transcripts upregulated in *p53*-KO cells compared to DKO cells revealed gene signatures relating to ribosome biogenesis and RNA biology, suggesting additional important aspects of the p53 pathway, to be explored in the future (Supplementary Fig. 5D).

To narrow the focus from thousands of differentially expressed transcripts called by RNA-seq to the most critical cancer driving pathways, we performed shotgun proteomics analyses on cells of each genotype. Validating the data set, we found that functional annotation of proteins downregulated in *p53-*KO cells compared to control cells indicated reduced expression of p53 target gene-encoded proteins, including APAF1, BAX, SESN2, and ZMAT3 (Supplementary Fig. 5E, Supplementary Table 9,10). We observed considerable overlap between proteins upregulated and downregulated in both DKO and *p53*-KO cells compared to control cells (Fig. 5C). Functional annotation of proteins more highly expressed in DKO and *p53-*KO cells relative to control cells again highlighted top biological categories related to cell migration, ECM, and integrin signaling (Fig. 5F-H, Supplementary Table 10). In addition, expression of other genes involved in cell division and cytoskeletal organization, such as CENPF and MAP2, were differentially expressed relative to control cells in DKO and *p53*-KO but not *Zmat3*-KO cells, suggesting cooperation between ZMAT3 and p21 in regulating these pathways (Fig. 5H, Supplementary Fig. 5F). Taken together, these analyses suggest that coordinated dysregulation of cell division, ECM, and cell migration programs are triggered by combined loss of p21 and ZMAT3 expression and are important consequences of loss of p53 signaling.

### Combined *Zmat3* and *Cdkn1a*-KO phenocopies *p53*-KO in enhancing cell migration

Because both transcriptomic and proteomic analysis of *Zmat3*-KO and DKO cells showed signs of dysregulated pathways involved in ECM organization and cell migration that overlapped with those identified in *p53*-KO cells, we sought to investigate the functional consequences of these gene signatures. To this end, we analyzed control, *Cdkn1a-*KO, *Zmat3-*KO, DKO, and *p53-* KO cells in a highly quantitative, 3D migration assay (34). In this assay, cells are embedded in collagen-I hydrogels and the ability of individual cells stained with octadecyl rhodamine B chloride to migrate through this matrix is examined by live-cell time-lapse confocal microscopy over the course of 24 hours (Fig. 6A), through measurement of both track length and mean speed for each cell. We noted first that *Cdkn1a*-KO and *Zmat3*-KO each caused a small but significant increase in track length and mean speed relative to control cells, suggesting that each gene individually mildly modulates cell motility (Fig. 6B-D). Interestingly, DKO cells showed clear enhancement of migration compared to either single knockout cell population, indicating that p21 and ZMAT3 cooperate to restrict motility (Fig. 6C-D). *p53-*KO also dramatically increased migration relative to wild-type control and single *Cdkn1a*-KO and *Zmat3*-KO cells, with an effect on track length similar to DKO cells but with a faster track mean speed than DKO cells (Fig. 6C-D). These results reveal that *Cdkn1a* and *Zmat3* combined knockout cooperatively increase migration to levels approaching those seen with p53 deficiency (Fig. 7). Together, these findings demonstrate that genetic cooperation between *Cdkn1a* and *Zmat3* DKO recapitulates important cancer-associated phenotypes of p53 deficiency, proliferation and migration, supporting their status as central effectors of p53-mediated tumor suppression.

**Figure 6.**
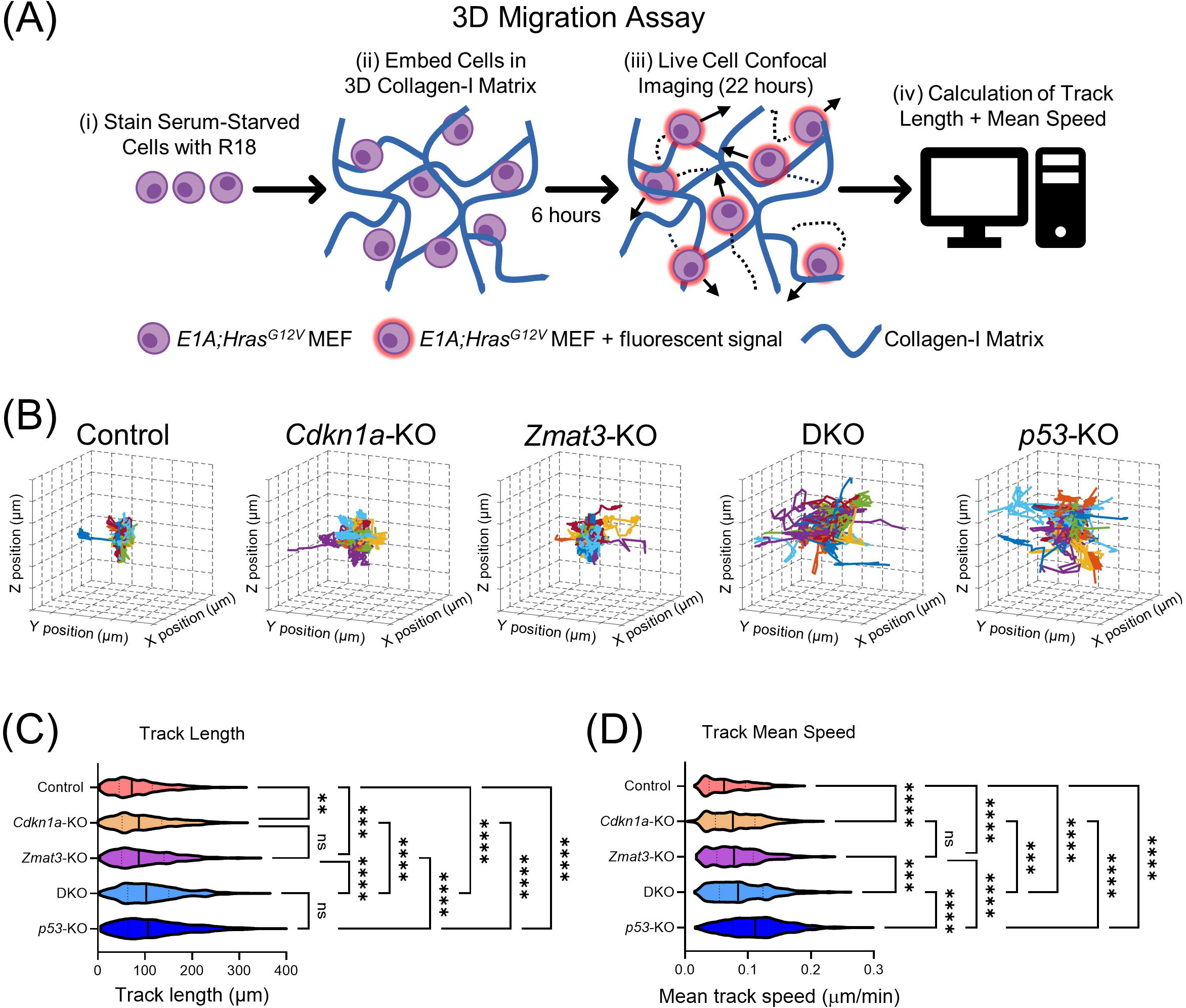
ZMAT3 and p21 deficiency cooperatively enhances cell migration in 3D, similar to p53 loss. **(A)** Schematic of 3D migration assay. (i) Serum starved *E1A;Hras^G12V^*MEFs are stained with octadecyl rhodamine B chloride (R18), (ii) embedded in a 3D collagen-I matrix, and, after 6 hours, are then tracked by live cell confocal microscopy for 22 hours. (iv) Track length and mean track speed are then calculated for individual cells. **(B)** 80 randomly selected cell trajectories shown for 3D migration experiments in *E1A;Hras^G12V^;H11^Cas9^* MEFs during a ∼22 hour period. Grid size = 20 µm. Different colors represent the tracks for different individual cells. **(C)** Track length (µm) and **(D)** mean speed (µm/min) for 3D migration experiments. (Control n = 746 cells, *Cdkn1a-*KO n = 701 cells, *Zmat3-*KO n = 659 cells, DKO n = 912 cells, *p53-*KO n = 833 cells). **p < 0.01, ***p < 0.001, ****p < 0.0001. Significance was calculated by ordinary one-way ANOVA with Tukey’s multiple comparisons test.

**Figure 7.**
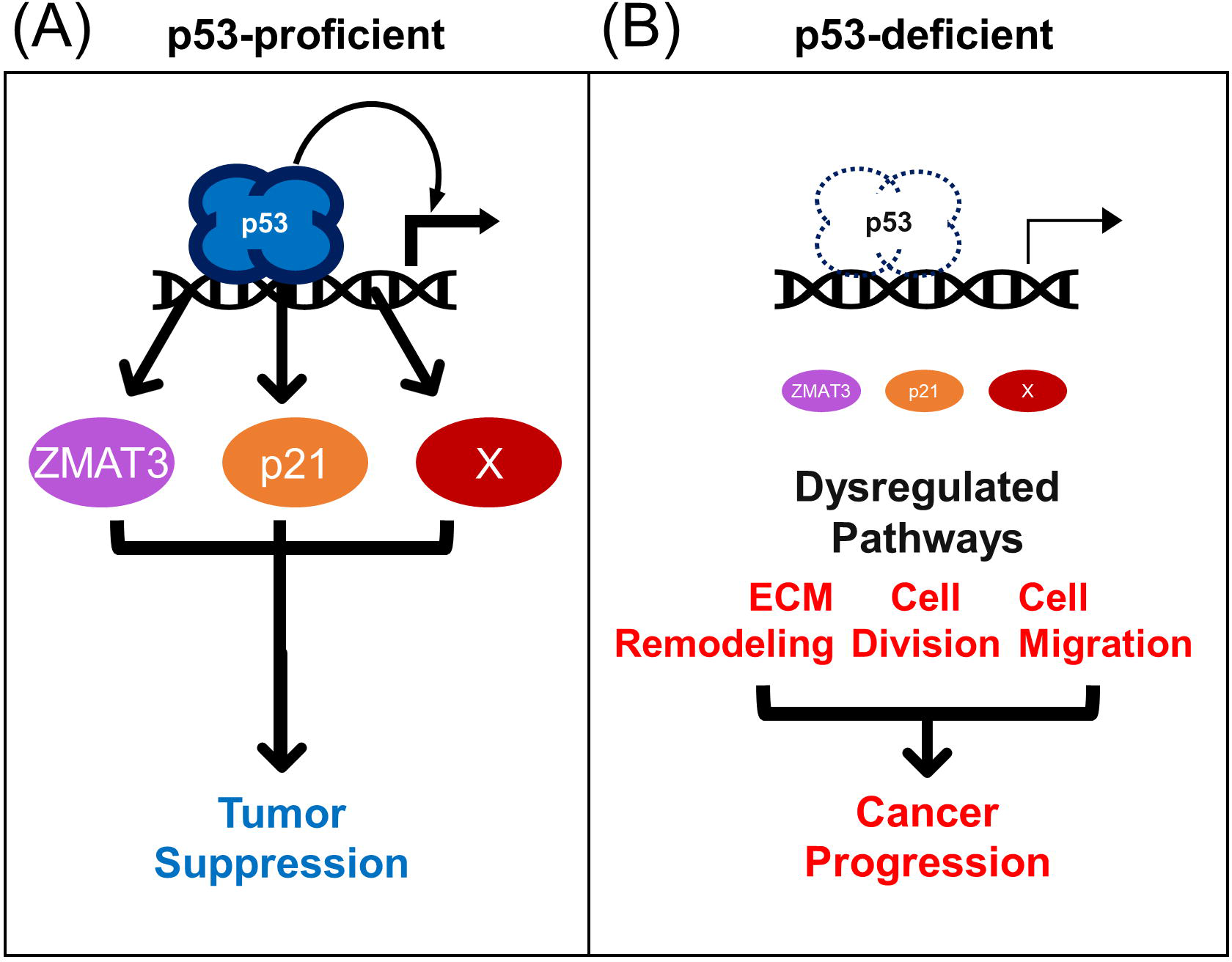
ZMAT3 and p21 are key effectors of p53-mediated tumor suppression. **(A)** Schematic for cooperative tumor suppression between *Zmat3* and *Cdkn1a* downstream of p53. X denotes components that might cooperate with ZMAT3 and p21. **(B)** In p53-deficient settings, transcriptional activation of target genes, like *Zmat3*, *Cdkn1a*, and X, diminishes, and causes changes to cellular behavior that promote cancer progression.

## Discussion

Despite the unequivocal importance of the tumor suppressor p53 in opposing tumorigenesis, our understanding of the mechanisms of p53-mediated tumor suppression remains incomplete. Here we use Tuba-seq^Ultra^ to shed new light on the strict p53-dependence of *Zmat3* in tumor suppression as well as on the relative contribution of the p53-*Zmat3* tumor suppression axis to p53-mediated tumor suppression as a whole, revealing that ZMAT3 accounts for roughly one third of the p53 tumor suppressor activity in initiation and growth, based on size and number. This finding is particularly striking given speculation that p53 distributes its tumor suppression capacity wide across its target gene network, with no individual target gene playing a significant role (35). Next, we used unbiased, combinatorial *in vivo* CRISPR/Cas9 screens to identify p53 target genes that cooperate with *Zmat3* in tumor suppression, revealing *Cdkn1a* as a top cooperator. Furthermore, through combined meta-analyses of p53-dependent gene expression in mouse and human data sets and of CRISPR/Cas9 knockout scores in human cells from Cancer Dependency Map data, we identify *ZMAT3* and *CDKN1A* as near universal p53 target genes that mediate tumor suppression in diverse cell types with varied driver mutations and across evolution. Integrated RNA-seq and proteomic analyses suggest that *Zmat3* and *Cdkn1a* knockout dysregulate a variety of phenotypes, including cell division and cell migration, which are also characteristic of p53-deficient cells. Consistent with these signatures, combined *Zmat3* and *Cdkn1a* knockout resulted in a dramatic change in cell behavior, as manifested in increased proliferation as well as enhanced cellular migration in a 3D matrix compared to control cells, akin to *p53* deletion. Thus, combined *Zmat3* and *Cdkn1a* loss recapitulates a significant part of p53 deficiency.

Consistent with our model highlighting the importance of *Zmat3* and *Cdkn1a*, a recent study reported the incidence of spontaneous cancer development in *Zmat3*^-/-^, *Bbc3^-/-^*, and *Cdkn1a^-/-^* compound mutant mice (36). Both *Cdkn1a*^-/-^;*Zmat3*^-/-^ and *Bbc3*^-/-^;*Zmat3*^-/-^ mice developed tumors, and *Bbc3*^-/-^;*Cdkn1a*^-/-^;*Zmat3*^-/-^ triple knockout (TKO) mice displayed significantly enhanced predisposition to spontaneous cancer development compared to either double knockout line or to wild-type mice, suggesting cooperation among all three targets in p53-mediated tumor suppression. Interestingly, a previous study of spontaneous tumorigenesis in *Bbc3*^-/-^;*Cdkn1a*^-/-^;*Pmaip1*^-/-^ mice from the same laboratory revealed no tumors through 500 days of aging (37), thus highlighting *Zmat3* inactivation as a critical element of tumorigenesis and reaffirming an essential role for *Zmat3* in cooperative tumor suppression. TKO mice did not exhibit as dramatic a tumor predisposition as that observed in *p53*-KO mice, supporting the notion that other p53 target genes must have tumor suppressor activity.

Although we found that *Zmat3* and *Cdkn1a* represent a significant portion of p53-mediated tumor suppression, our data also suggest that p53 mediates tumor suppression through additional target genes. The fact that we did not identify other widespread tumor suppressor target genes through our approaches could be explained by those genes serving more redundant functions (35) or highly tissue-specific roles (38), both of which would fail to score dramatically in our assays. An important clue for tumor suppressive functions of p53 outside of those regulated by *Zmat3* and *Cdkn1a* has come from molecular comparison of DKO and *p53-*KO cells, which uncovered ribosome biogenesis and RNA homeostasis signatures upregulated with *p53*-KO relative to DKO gene expression. Indeed, p53 serves as a key repressor of translation (39,40) and numerous studies have linked p53 to inhibition of mTOR, a complex that can promote protein synthesis (41–43). It is possible that mTOR inhibition might be widely important for p53 tumor suppressor function across contexts, but may not have scored in our assays or in our DepMap analysis because there are numerous p53-inducible inhibitors of mTOR, such as *SESN1* and *SESN2* (44), that are either tissue-specific or redundant with each other. In addition, our study focused on protein-coding p53 target genes, but p53-induced non-coding RNAs represent additional potential effectors of p53-mediated tumor suppression (45).

At a molecular level, ZMAT3 loss dysregulates expression of genes involved in ECM organization and cell migration at both the RNA and protein level and, furthermore, increases cell migration relative to control cells. Inspection of individual differentially expressed genes in ZMAT3 deficient cells reveals potential therapeutic targets such as ITGA3 and ITGA6, which are upregulated in *Zmat3*-KO, DKO, and *p53-*KO cells relative to control cells at both the RNA and protein levels and are associated with poor patient prognosis in the clinic and aggressive cellular behavior in culture (46,47). This enhanced cell motility with ZMAT3 loss, which is further enhanced by p21 inactivation, likely reflects fundamental changes in cellular architecture, such as in the cytoskeleton, which may affect cell behavior in various ways.

Our study addresses the need for better understanding the mechanisms of p53-mediated tumor suppression by advancing identification of signaling pathways and cellular processes that might be broadly targeted by cancer therapy in p53-deficient tumors. Genes upregulated with combined *Cdkn1a* and *Zmat3* loss, or with p53 inactivation, such as *Itga3*, encoding an integrin subunit, are of particular interest for therapeutic inhibition. Integrins have long been considered prime targets for anti-cancer therapies (46). Future work will need to investigate which proteins we have identified are most suitable for clinical development based on functional oncogenic roles and our ability to target them therapeutically. Ultimately, by deconstructing the p53 pathway, we take one step closer to targeting the p53 pathway in the clinic.

## Materials and Methods

### Cell culture

Mouse embryonic fibroblasts were maintained in Dulbecco’s Modified Eagle Medium (Gibco) supplemented with 10% Fetal Calf Serum (FCS) and incubated at 37°C in a carbon dioxide incubator held at 5% oxygen. Doxorubicin (Sigma, Cat # D1515) treatment was done at a concentration of 0.2 μg/ml for 8 hours. *E1A;Hras*^G12V^;*H11^Cas9^* MEFs were generated as previously described(3,15). 293AH viral packaging cells were maintained in Dulbecco’s Modified Eagle Medium (Gibco) supplemented with 10% Fetal Calf Serum (FCS) and incubated at 37°C in a carbon dioxide incubator held at 20% oxygen. 293AH cells were transduced by Lipofectamine™ 2000 (Thermo Fischer Scientific) in Opti-MEM™ (Thermo Fischer Scientific) according to manufacturer’s instructions with viral packaging agents VSV-G and Δ8.2 plus lentiviral expression plasmid. Puromycin (2 μg/ml) and Neomycin (550 ug/mL) were used to select for target cells that had stably integrated lentiviral expression plasmids with the cognate resistance markers. The sex of the mouse cell lines was not determined as it was not expected to impact the results. Cell line authentication was not applicable.

### *Zmat3* conditional knockout mouse line generation

The *Zmat3* conditional knockout allele (*Zmatf3^fl^*) was designed to insert loxP sites at *Zmat3* intron 3 and intron 5, floxing exon 4 and exon 5. Deletion of exon 4 and exon 5 after Cre expression was predicted to result in a downstream reading frame shift and 159 amino acid deletion. *Zmat3^fl^* mice were generated using CRISPR/Cas9-mediated genome recombination. Five guide RNAs were designed, and sgRNA were purchased from Synthego (Menlo Park CA). After *in vitro* validation by DNA cutting efficiency, two sgRNAs were chosen for generating *Zmat3^fl^* mice, *Zmat3*-g3 (5’-CTAGTATCCGTTGAGAAAAT-3’) and *Zmat3*-g5 (5’-CACCCACTACCTAATTCCCG-3’). The 2426bp donor DNA contained 1000bp homologous arms on each side. The donor DNA was cloned into a vector plasmid. Double strand donor DNA fragments were extracted from the plasmid vector after restriction enzyme digestion. A mixture of sgRNA (16ng/ul), Cas9 protein (25ng/ul, IDT) and donor DNA (3ng/ul) were injected into C57Bl/6J mouse zygote pronuclei. 271 embryos were injected, 56 pups were born, and 6 pups were identified by PCR and DNA sequencing. The genotyping PCR confirming 5’ loxP insertion into the targeted region of the *Zmat3* locus consisted of one external primer upstream of donor DNA (5’-CAGGTCCACATGTCGTGCATAC-3’) and one primer overlapping the 5’ loxP site (5’-CAACTCAGGAAAGCACATAACTTCG-3’). The PCR product was checked by sequencing. The same strategy confirmed proper integration of the 3’ loxP site into the intended lcous using one external primer downstream of the donor DNA (5’-CTGTGCCATCCCCATCCTAAATG-3’) and one primer at the 3’ loxP site (5’-GTCCACACTTTAAACCCATTATAACTTCG-3’). The *Zmat3^fl^*founder mice were bred with C57Bl/6J pure background mice and then crossed with appropriate additional mice for experimentation. A secondary genotyping PCR was developed using the following primers (see Supplementary Fig. 1):

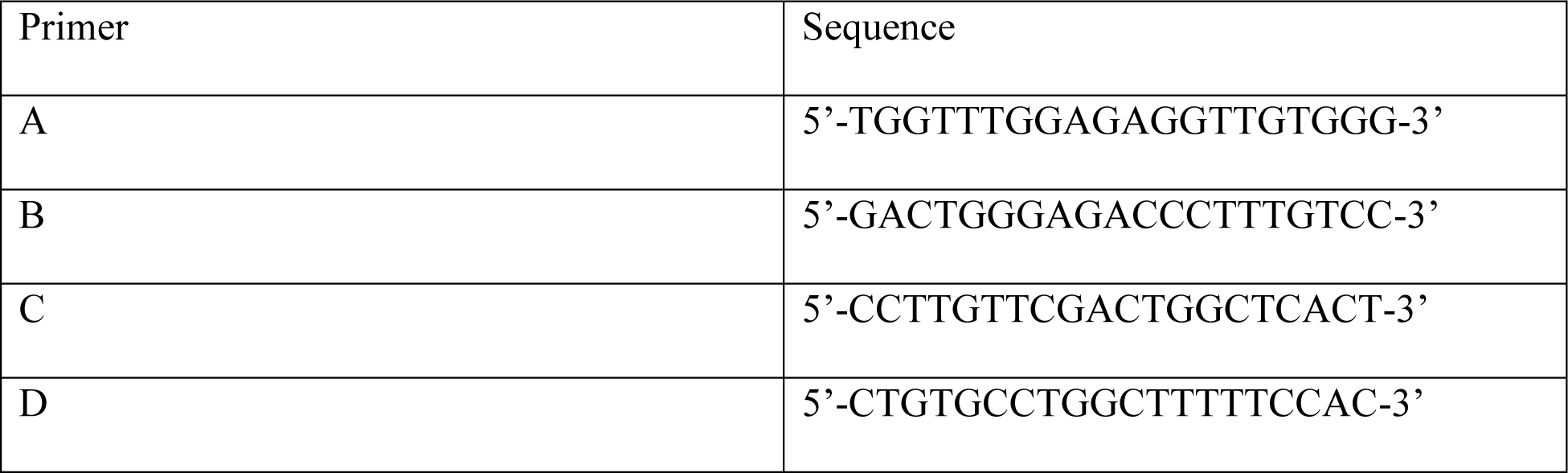

### Tumor barcode sequencing

A U6-integrated diverse barcode-sgRNA/Cre library containing sg*Trp53*, sg*Zmat3*, sg*Zmat3* RE, and sgInerts was generated, packaged into lentivirus and titered as previously described (21). Briefly, the lentiviral vector expressing sgRNA and Cre recombinase (Tuba-seq^Ultra^) vector backbone was barcoded by Gibson assembly. The sgRNA pool was synthesized on oligo chips (Twist Biosciences, CA) and cloned into the barcoded vector via Golden Gate assembly. The assembled plasmid library was electroporated into 10-beta electrocompetent *E. coli* (NEB, C3020K) as described by the manufacturer. The bacterial colonies were pooled, and the plasmids were extracted using Qiagen Midi-prep (Qiagen, #12941). Lentiviral packaging was performed by co-transfecting library plasmids with pCMV601 VSV-G (Addgene, 8454) envelope plasmid and pCMV-dR8.2 dvpr (Addgene, 8455) packaging plasmid using Opti-MEM (Gibco, 31985070) in 150 mm cell culture plates. The virus was concentrated by ultracentrifugation (25,000g for 1.5 hours at 4°C), and resuspended in 1X PBS.

Mice aged 8 weeks or older of *KT*, *KT;H11^Cas9^*, *KZT;H11^Cas9^* or *KPT;H11^Cas9^* genotype were anesthetized by intraperitoneal injection of Avertin (2-2-2 Tribromoethanol) and infected by intratracheal instillation with 300k, 150k, 100k, and 30k ifu lentivirus packaged using the U6-integrated diverse barcode-sgRNA/Cre library respectively. Mouse lungs were collected 15 weeks after intratracheal infection, weighed, and frozen for further processing.

Genomic DNA was isolated from bulk tumor-bearing lung tissue from each mouse, and benchmark control cell lines were added to each mouse lung sample prior to lysis to enable the calculation of the absolute number of neoplastic cells in each tumor from the number of barcode-sgRNA reads as previously described (21). The genomic DNA were prepared for next-generation sequencing (NGS) by amplifying the barcode-sgRNA region using high-fidelity DNA polymerase (NEB, M0544S) with primers compatible for Illumina sequencing platforms (Tang et al., 2024). The NGS library was sequenced with 150bp pair-ends on Novaseq 6000 by Novogene (Novogene, CA).

Paired-end reads were first merged using AdapterRemoval (48), and merged reads were parsed using regular expressions to identify the sgRNA sequence and clonal barcode. For identifying sgRNA sequences, we required a perfect match with the designed sequences, as mutated sgRNA could suffer from reduced efficiency and off-target effects. The 12-nucleotide clonal barcode possesses a high theoretical diversity of clonal barcode (> 10^6^) ensuring each clonal tumor is uniquely barcoded. As the probability of two clonal tumors with the same tumor genotype possessing two barcodes within a 1-hamming distance of each other is extremely low, when we encountered low-frequency clonal barcodes within a 1-hamming distance of high-frequency clonal barcodes, we attributed them to sequencing or PCR errors (spurious reads). These low-frequency barcodes were merged with barcodes of higher frequencies. Finally, total reads for each genotype-barcode combination were calculated. Next, we converted the read counts associated with clonal tumors into absolute neoplastic cell numbers. This conversion was accomplished by normalizing the reads of the clonal tumor to the number of reads of the “spike-in” benchmark cell lines added to each sample prior to lung lysis and DNA extraction. We imposed a minimum tumor size cutoff of 300 cells for downstream analysis, which ensured at least two reads for the smallest clonal tumor.

To quantify the impact of each gene on tumor growth, we normalized tumor statistics for a given sgRNA *X* (sg*X* tumors) against control sgRNAs (sgInert tumors). We used two key measures: the size of tumors at defined percentiles of the distribution (specifically the 50^th^, 70^th^, 80^th^, 90^th^, and 95^th^ percentile tumor sizes), and the log-normal mean (LN mean) size. The LN mean represents the average tumor size, assuming a log-normal distribution. By normalizing these measures to the corresponding sgInert statistics, we derived ratios that indicate the growth advantage or disadvantage of each tumor genotype compared to sgInert tumors.

When calculating the log-normal mean with an adaptive sampling method, rather than using the same cell number cutoff (300 cells) for all tumors, we applied varying size cutoffs for tumors associated with different sgRNAs. This approach ensured that the tumor number ratios between a focal sgRNA and sgInert were consistent across *KT;H11^Cas9^, KZT;H11^Cas9^*, and *KPT;H11^Cas9^* mice, as they were in *KT* mice.

In addition to the tumor size metrics, we characterized the effects of gene inactivation on tumorigenesis by evaluating the number of tumors (“tumor number”) associated with each genotype. To assess the extent to which a given gene (*X)* affects tumor number, we therefore first normalized the number of sg*X* tumors in *KT;H11^Cas9^* mice (also in *KZT;H11^Cas9^* and *KPT;H11^Cas9^*) by the number of sg*X* tumors in the *KT* mice. As with the tumor size metrics, we then calculated a relative tumor number by normalizing this statistic to the corresponding statistic calculated using sgInert tumors. Analogous to the calculation of relative tumor number, we characterized the effect of each gene on tumor burden by first normalizing the sgX tumor burden in Cas9-expressing mice to the burden in *KT* mice. We then calculated a relative tumor burden by normalizing this number to the corresponding statistic calculated using sgInert tumors. The confidence intervals and *P*-values were calculated using a bootstrap resampling approaches with 10000 cycles, as previously described (20). To account for multiple hypothesis testing, *P*-values were FDR-adjusted using the Benjamini-Hochberg procedure (49).

The gene-level metrics (M_gene_) were estimated using a weighted average of sgRNA-level metrics (M_sgRNA_):

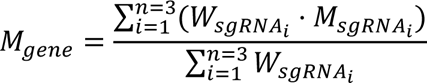

where *W_sgRNA_i__* is the relative tumor number of sgRNA_i_ for the focal gene.

### sgRNA-expression plasmid cloning

GH020 (see Addgene_85405) and MCB320 (see Addgene_89359) lentiviral expression plasmids were used to express sgRNA. sgRNA were cloned into plasmids by restriction enzyme digest method (GH020: BbsI-HF®, MCB320: BlpI+BstXI) and propagated in Subcloning Efficiency™ DH5α Competent Cells (Thermo Fischer Scientific) selected with Ampicillin (100 μg/mL). The following table relates the sgRNA sequences used:

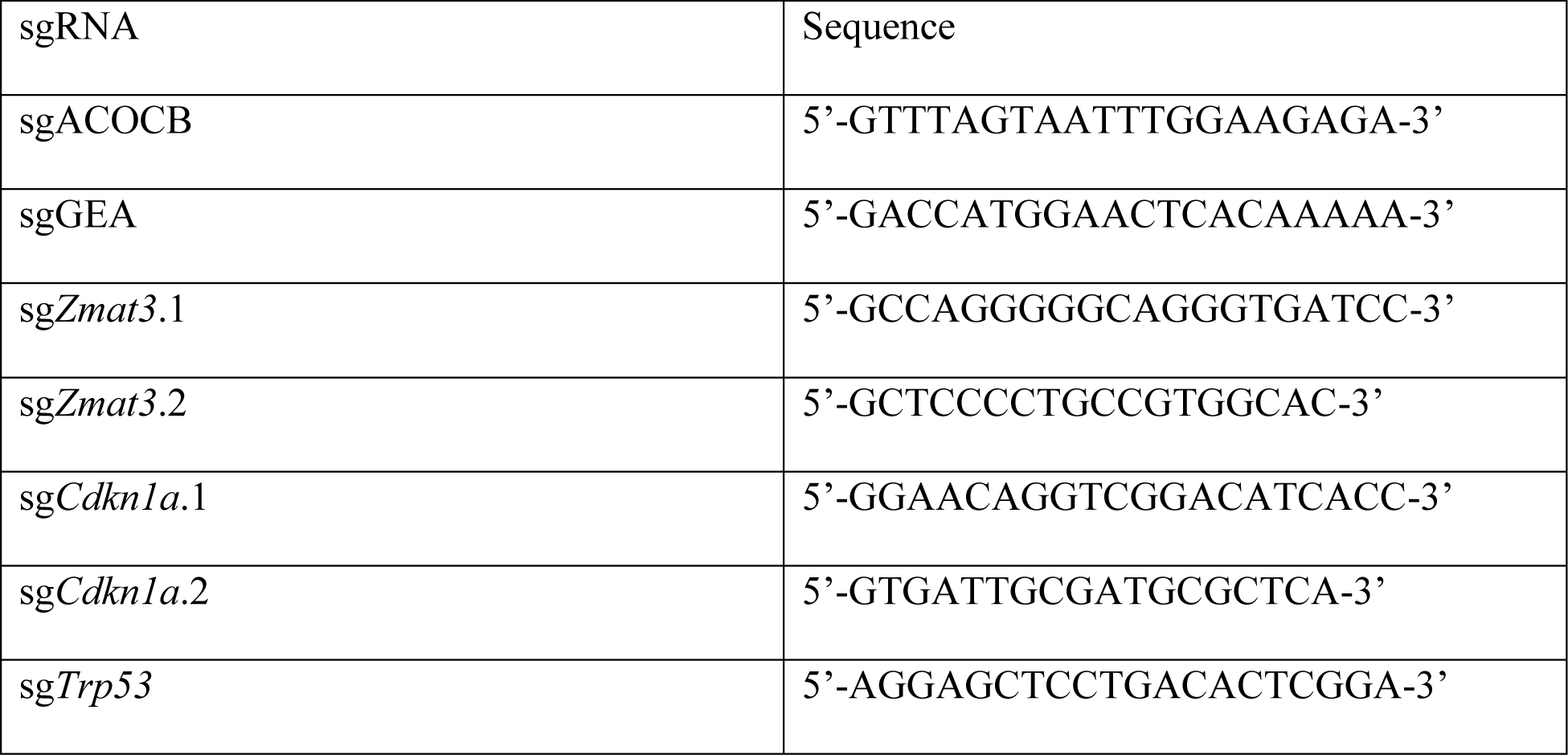

### Flow cytometry

Cells were analyzed for GFP and mCherry expression by flow cytometry (LSR Fortessa, BD Biosciences).

### qRT-PCR

RNA extraction was performed using TRIzol reagent (Thermo Fisher Scientific, Cat #15596018) according to the manufacturer’s protocol. Reverse transcription was conducted with M-MLV reverse transcriptase (Thermo Fisher Scientific, Cat # 28025) and random primers (Thermo Fisher Scientific, Cat # 48190). 1 μg of total RNA was used for cDNA synthesis. cDNA was diluted 1:5 in nuclease-free water and stored at –80°C until used. Quantitative PCR was performed in triplicate using PowerUP SYBR green master mix (Thermo Fisher Scientific, Cat # A25743) and a 7900HT Fast Real-Time PCR machine (Applied Biosystems). Expression analysis was performed using the following primers: *Cdkn1a* (5′-CACAGCTCAGTGGACTGGAA-3′, 5′-ACCCTAGACCCACAATGCAG-3′), *Mdm2* (5’-CTGTGTCTACCGAGGGTGCT-3’, 5’-CGCTCCAACGGACTTTAACA-3’), *Gapdh* (5’-AGGTCGGTGTGAACGGATTTG-3’, 5’-TGTAGACCATGTAGTTGAGGTCA-3’ The mean of the housekeeping gene *Gapdh* was used as an internal control to normalize the variability in expression levels. All qRT-PCR performed using PowerUP SYBR Green was conducted at 50°C for 2 min, 95°C for 10 min, and then 40 cycles of 95°C for 15 s and 60°C for 1 min. Melt curve analysis was done to verify the specificity of the primers. Samples were quantified using a standard curve method.

### Western blot

Western blots were performed according to standard protocols. Briefly, cells were lysed in RIPA buffer (50 mM Tris pH 8.0, 150 mM NaCl, 1% NP-40, 0.5% sodium Deoxycholate, 0.1% SDS) or SDS extraction buffer (2% SDS, 50 mM Tris pH 6.8) with freshly added Complete protease Inhibitor Cocktail (Roche). Extracts were run on 10% polyacrylamide SDS-PAGE gels, gels were transferred to PVDF membrane (Immobilon, Millipore), and membranes were blocked with 5% milk in TBST and probed with antibodies directed against ZMAT3 (1:1000, Santa Cruz), p53 (CM5, 1:1000 Leica Novocastra), p21 (EPR18021, 1:1000, abcam) or GAPDH (1:20,000, Fitzgerald), followed by anti-mouse or anti-rabbit HRP-conjugated secondary antibodies (1:5,000, Vector Laboratories). Blots were developed with ECL Prime (Amersham) and imaged using a ChemiDoc XRS+ (BioRad).

### In vivo screen

All animal experiments were in accordance with the Stanford University APLAC (Administrative Panel on Laboratory Animal Care). The p53 target gene sgRNA library was synthesized and cloned into MCB320 lentiviral expression vector as previously described, with 10 sgRNA for each of the 272 p53 target genes and 1000 safe-harbor sgRNAs, which target predicted non-functional regions in the mouse genome to replicate the effects of DNA damage induced by Cas9(50,51). For the WT screen, the lentiviral p53 target gene sgRNA library was transfected into 293AH cells with TransIT®-LT1 Transfection Reagent (Mirus, MIR2304). 48 hours later lentivirus was transduced into target cells, followed by selection with puromycin as previously described(15). The *Zmat3*-KO and *p53*-KO screen cells were simultaneously infected with the lentiviral p53 target gene sgRNA library and GH020-sg*Zmat3*.2 or GH020-sg*Trp53*. These cells went through double selection with puromycin and neomycin to select for the sgRNA library and the GH020-sgRNA plasmid, respectively. Libraries were designed, subcutaneous tumors were formed, genomic DNA was harvested and prepped for sequencing, and casTLE analysis was performed as previously described(15) except that the screens were quantified as two replicates (R1 and R2), each comprising genomic DNA from multiple tumors. Briefly, for each screen, 4-5 mice were engrafted with 3 million cells mixed with 200µL of 50:50 Matrigel®(Corning):PBS by volume on both flanks of 10-12-week-old male ICR/Scid mice (Taconic). Tumors were harvested after three weeks of growth. Frozen tumors were ground in liquid nitrogen, followed by genomic DNA preparation (Gentra Puregene). Genomic DNA preparations extracted from the left flank tumors and the right flank tumors were pooled respectively by mixing equal amounts giving n = 2 replicates, R1 and R2. MEMcrispR(29) was carried out on the counts from the screen in addition to casTLE with the modification that the T0 data were run in duplicate for each of the 3 analyses. ICR/Scid mice were group-housed (up to 5 mice per cage), and irradiated chow and water were provided *ad libitum*.

### RNA-sequencing and analysis

5 genotypes of cells were generated in *E1A;Hras*^G12V^;*H11^Cas9^*MEFs. Each line was iteratively infected and selected with lentivirus to deliver GH020 plasmid (GFP, neomycin selection) and then MCB320 plasmid (mCherry, puromycin). The following table shows the sgRNA delivery strategy:

| Genotype | Replicate | Plasmid 1 | Plasmid 2 |
| --- | --- | --- | --- |
| Control | 1 | GH020-sgACOCB | MCB320-sgGEA |
| Control | 2 | GH020-sgGEA | MCB320-sgACOCB |
| *Cdkn1a*-KO | 1 | GH020-sgACOCB | MCB320-sg*Cdkn1a*.1 |
| *Cdkn1a*-KO | 2 | GH020-sgACOCB | MCB320-sg*Cdkn1a*.2 |
| *Cdkn1a*-KO | 3 | GH020-sgGEA | MCB320-sg*Cdkn1a*.1 |
| *Cdkn1a*-KO | 4 | GH020-sgGEA | MCB320-sg*Cdkn1a*.2 |
| *Zmat3*-KO | 1 | GH020-sg*Zmat3*.1 | MCB320-sgACOCB |
| *Zmat3*-KO | 2 | GH020-sg*Zmat3*.1 | MCB320-sgGEA |
| *Zmat3*-KO | 3 | GH020-sg*Zmat3*.2 | MCB320-sgACOCB |
| *Zmat3*-KO | 4 | GH020-sg*Zmat3*.2 | MCB320-sgGEA |
| DKO | 1 | GH020-sg*Zmat3*.1 | MCB320-sg*Cdkn1a*.1 |
| DKO | 2 | GH020-sg*Zmat3*.1 | MCB320-sg*Cdkn1a*.2 |
| DKO | 3 | GH020-sg*Zmat3*.2 | MCB320-sg*Cdkn1a*.1 |
| DKO | 4 | GH020-sg*Zmat3*.2 | MCB320-sg*Cdkn1a*.2 |
| *p53*-KO | 1 | GH020-sg*Trp53* | MCB320-sgACOCB |
| *p53*-KO | 2 | GH020-sg*Trp53* | MCB320-sgGEA |

To generate n = 4 for Control cells and *p53*-KO cells, cells were plated in duplicate from each of the two lines. RNA was extracted from cells using the RNeasy midi kit (Qiagen, Cat#75144). RNA quality was assessed using a BioAnalyzer (2100 Bioanalyzer Instrument, RRID:SCR_018043). Novogene generated and sequenced RNA-seq libraries using the Illumina TruSeq RNA Sample Preparation kit (Cat# RS-122-2001) and Illumina HiSeq4000 (RRID:SCR_016386). Reads were aligned to the GRCm38 mouse genome using HISAT2 (v 2.0.4; RRID:SCR_015530). Sorted BAM files were generated using Samtools (v 1.3.1; RRID:SCR_002105). The number of reads mapping to each gene in the mouse Ensembl database (v 87; RRID:SCR_002344) was counted using HTSeq-count (v 0.6.1; RRID:SCR_011867). Differential gene expression analysis was performed using DESeq2 (v 1.24.0; RRID:SCR_015687). Functional annotation was performed using Metascape (52).

### Shotgun proteomics

4 genotypes of cells were generated in *E1A;Hras*^G12V^;*H11^Cas9^*MEFs. The following table shows the sgRNA delivery strategy:

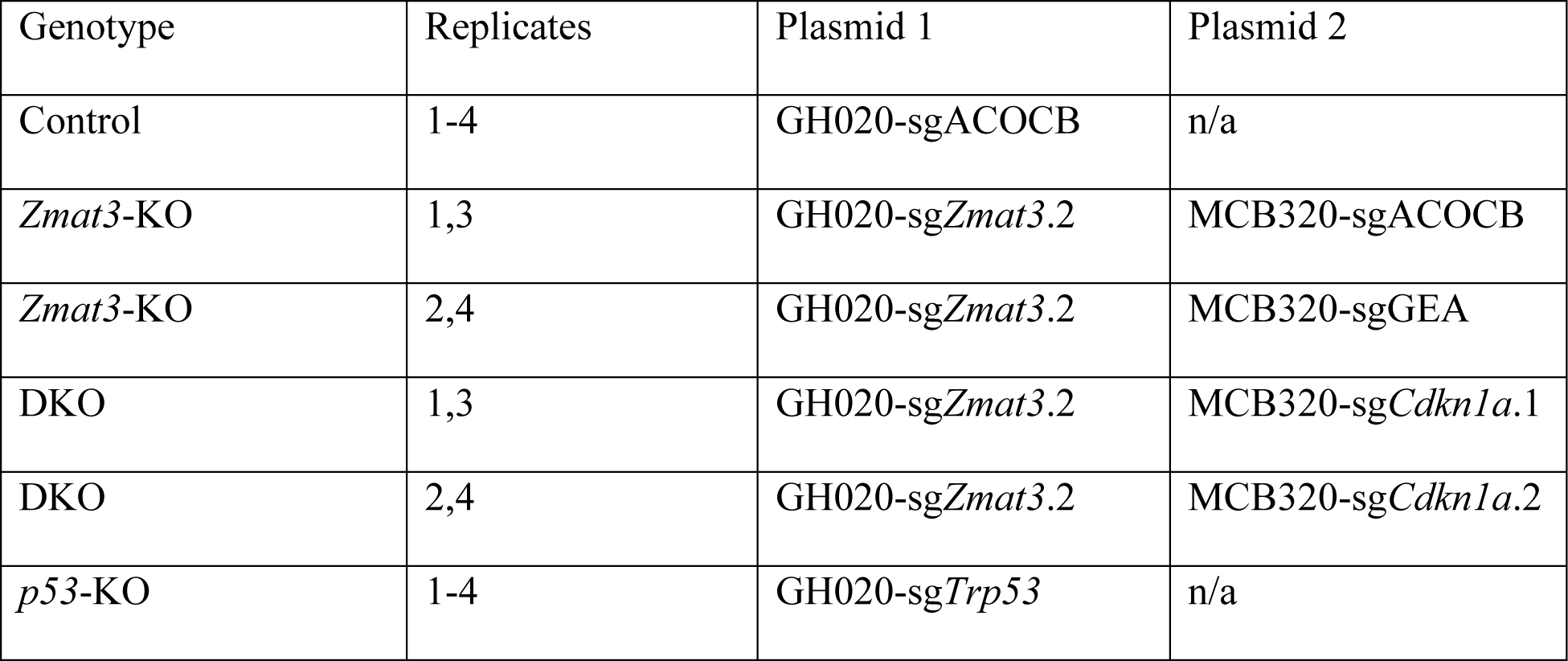

Cells were either plated in quadruplicate or in duplicate to generate n = 4 for each genotype. Cell lines were lysed in the plate and then protein was extracted using the iST 8x kit (Preomics P.O.00001) according to the manufacturer’s instructions. Prepared samples were dried using a SpeedVac and resuspended in Solution A (2% ACN, 0.1% FA). Sample concentrations were quantified using Quantitative Colorimetric Peptide Assay (ThermoFischer, 23275) and diluted to 100 ng/µl. 200 ng of samples were analyzed using the timsTOF Pro (Bruker Daltonics). Liquid chromatography was performed using PepSep Ten Columns (I.D. 75 µM, particle size 1.9 µm, Bruker Daltonics, 1893472) at 50°C and peptides were separated using a 68 min gradient (solvent A: 2% ACN, 0.1% FA; solvent B: 0.1% FA in ACN) at a flow rate of 500 nl/min. A linear gradient from 3-33% B was applied for 60 min, followed by a step to 95% B for 5 min and 3 min of washing at 95% B. The timsTOF Pro was operated in PASEF mode with the following settings: Mass Range 100 to 1700m/z, 1/K0 Start 0.85 V·s/cm2, End 1.3 V·s/cm2, Ramp time 100ms, Lock Duty Cycle to 100%, Capillary Voltage 1700, Dry Gas 3 l/min, Dry Temp 200°C, PASEF settings: 4 MS/MS, charge range 0-5, active exclusion for 0.04 min, Scheduling Target intensity 20000, Intensity threshold 500, and CID collision energy 10eV.

Bruker raw data files were searched against a database of NCBI mouse proteome (32,687 entries, updated 20 July 2021, only reviewed proteins) supplemented with common contaminants and reverse decoy sequences using MSFragger (version 3.5)(53) run via FragPipe (v18.0)(54). Mass tolerances on precursors and fragments were ≤20 ppm and ≤40 ppm, respectively; protein cleavage was set to “specific” with a maximum of 2 missed cleavages per peptide. Data type was DDA, and the enzyme was trypsin, cleaving after residue K and R but not before P. Carbamidomethylation (C) was specified as fixed while oxidation (M), acetylation (protein N-terminal) and Ser/Thr/Tyr phosphorylation were set as variable modifications. Peptide-, PTM- and ProteinProphet (55) were used for validation. Search results were then imported into Skyline (v22.1.9) (56) for ms1 quantitation using the same mass tolerances and modifications. Peak intensities were exported in MSstats format for differential analysis in R. Intensity results were summarized using MSstats(57) DataProcess function with censoredInt set to ‘0’, featureSubset to ‘highQuality’ and remove_uninformative_feature_outlier to T, with the other parameters left as default. The summarized protein level data was formatted for analysis with DEP package (58). Pathway analysis was performed using Metascape (v 3.5; RRID:SCR_016620) on proteins significantly upregulated or downregulated as called by padj < 0.05.

### Analysis of *TP53* mutation-specific *ZMAT3* and *CDKN1A* dependencies in DepMap

*ZMAT3* and *CDKN1A* genetic dependencies, in relation to *TP53* status, were assessed in using publicly-available data from the Cancer Dependency Map (DepMap). Somatic mutations, copy number status, CRISPR gene effect scores, gene expression, and cell line metadata files were downloaded from the DepMap Public 23Q4 Primary Files release. Cell lines were annotated as *TP53*^Deficient^ if they (1) contained variants annotated as CosmicHotspot or CCLEDeleterious, (2) had a *TP53* copy number status of < 0.5, or (3) contained multiple mutations or a combined mutation and deletion. In total, 1,087 cell lines contained gene effect and *TP53* mutational data and 1,450 cell lines contained gene expression and *TP53* mutational status.

Differential effect scores were calculated for all genes pan-cancer (n = 1,087). Scores were calculated by subtracting the mean effect score for all *TP53*^WT^ cell lines by the mean effect score for all *TP53*^Deficient^ cell lines. P-values and adjusted p-values were calculated for each comparison using the t.test() and p.adjust() functions in the R stats package (version 3.6.2). Genes were then rank-ordered by effect difference such that genes with a negative score were more dependent in *TP53*^WT^ cell lines and genes with a positive score were more dependent in *TP53*^Deficient^ cell lines. We further separated genes into two bins: those where the mean gene effect scores in both *TP53*^Deficient^ and *TP53*^WT^ were positive (suggesting tumor suppressor activity) and those where the mean gene effect scores in both *TP53*^Deficient^ and *TP53*^WT^ were negative (suggesting a synthetic lethal-like relationship). Finally, using a curated list of 343 known p53 target genes (32), we performed the same pan-cancer and cancer-specific analyses to determine *ZMAT3* and *CDKN1A* ranks.

Differential gene expression analysis between *TP53*^WT^ and *TP53*^Deficient^ cell lines was performed pan-cancer (n = 1,450). Differentially expressed genes were detected by first filtering for genes with at least five samples containing at least ten reads per gene and then applying DESeq2 (version 1.40.2) to the remaining unnormalized read counts. Genes with a log2 fold-change ≥ 1.5 and adjusted p-value < 0.01 were considered significantly upregulated in *TP53*^Deficient^ lines while those with a log2 fold-change ≤ 1.5 and adjusted p-value < 0.01 were considered significantly downregulated in *TP53*^Deficient^ lines.

All code required to download, process, and visualize these DepMap analyses is provided at https://github.com/brooksbenard/tp53_p21_zmat3/tree/main.

### Meta-analysis of p53 gene regulation and binding

Previously published data on p53-dependent gene regulation were retrieved from www.TargetGeneReg.org (31), which contains 15 mouse and 57 human transcription profiling datasets. Additionally, genome-wide data on p53 binding were available from 9 curated ChIP datasets derived from mouse cells (33) and 28 ChIP datasets from human cells (59).

### Proliferation assay

*E1A;Hras^G12V^;H11^Cas9^* MEFs were plated in duplicate in a 12-well plate at 30,000 cells per well. At 24, 48, 72, and 96 hours cells were harvested, centrifuged, resuspended in 1mL of DMEM, and counted by a Luna II Automated Cell Counter. Fold change in cell number was calculated with respect to t = 24 hours. Two replicates were performed in duplicate (total n = 4).

### 3D Migration in Collagen-I hydrogels and Image Analysis

For live-cell time-lapse imaging, *E1A;Hras*^G12V^;*H11^Cas9^* MEFs of different genotypes were first starved in serum-free medium overnight. Cell were trypsinized and resuspended in cell culture media. To visualize cells within the 3D hydrogel, cells were stained with octadecyl rhodamine B chloride (R18, 1:1,000 dilution, stock 10mg mL^-1^; Thermo Fisher Scientific) prior to 3D cell encapsulation.

To form cell-laden 3D collagen-I hydrogels, reconstituted collagen-I (TeloCol-10, Advanced Biomatrix) was first diluted to a final working concentration of 3 mg mL^-1^ in DPBS. The collagen-I solution was then neutralized to pH 7.0. R18 labeled cells were gently mixed into the hydrogel precursor solution at a final concentration of 1.5M mL^-1^. The cell-laden hydrogel precursor solution was then cast into a custom made 1.5 w/v % agarose (GibcoBRL 15510-027) gel molds within a µ-Slide 2 Well Imaging chamber (ibidi 80286) and then incubated at 37°C for 45 minutes. Fresh medium was added after hydrogel gelation. After 6 hr, encapsulated cells were imaged at 20 min intervals for ∼22 hr using a Leica STELLARIS 5 confocal microscope with HC PL APO 10x/0.40 air objective. 3D z-stacks (62.5 µm, with 12.5 µm z-intervals) were captured at each position. Cell-laden hydrogels were kept in an incubated chamber (37°C and 5% CO_2_) during the live-cell time-lapse experiments.

For cell migration analysis, Imaris software (Bitplane) was used to track R18 labeled cells from live-cell time-lapse imaging experiments. A Surface rendering analysis was used to track single cell migration that were captured for at least 17 hr during the live-cell experiment. Specific parameters were adapted as previously reported (34). Mean track speed and track length were automatically calculated by Imaris. To reconstruct 3D cell migration trajectories, a custom MATLAB script was used from the Imaris cell track position output.

## Competing Interests Statement

R.M. is on the Advisory Boards of Kodikaz Therapeutic Solutions, Orbital Therapeutics, Pheast Therapeutics, 858 Therapeutics, Prelude Therapeutics, Mubadala Capital, and Aculeus Therapeutics. R.M. is a co-founder and equity holder of Pheast Therapeutics, MyeloGene, and Orbital Therapeutics. The other authors declare no competing interest.

## Acknowledgements

We thank Julien Sage for critical reading of the manuscript and Sydney Lu, Scott Dixon, and Camila Bolle for important discussions. We thank Mingxin Gu for assistance with the *in vivo* screens. We thank Nitin Raj, Sofia Ferreira, and Kathryn Hanson for advice on RNA-sequencing. We thank Richard Frock for the use of his transilluminator and SpeedVac and Laura Andrejka for assistance with Tuba-seq^Ultra^. This work was supported by Tobacco-Related Disease Research Program (TRDRP) fellowship T31DT1713, NIH T32CA009302 to A.M.B, NIH-R01CA234349 (to D.A.P and M.M.W), and NIH R35CA197591 and TRDRP grant 28IP-0037 to L.D.A. We apologize to those whose work we could not cite due to spatial constraints.

## Author contributions

A.M.B. and L.D.A. designed experiments and interpreted the results. A.M.B. performed experiments and analyzed data. A.R.M. conducted experiments. D.Y. conducted casTLE analysis on sgRNA screens. H.X., M.W., Y.J.T., and S.S.L. performed Tuba-seq^Ultra^ pipeline. J.D. and R.C. processed and analyzed shotgun proteomics samples. L.J.V. designed the sgRNA library. B.A.B. performed analyses of DepMap data. S.S. performed and analyzed 3D migration experiments. M.F. conducted meta-analysis of p53-dependent gene expression and ChIP-seq binding. F.Y. helped interpret 3D migration experiments. R.M. helped interpret DepMap data. P.K.J. helped interpret shotgun proteomics data. D.A.P and M.M.W. helped interpret Tuba-seq^Ultra^ data. M.C.B. helped interpret sgRNA screen results. A.M.B. and L.D.A. wrote the manuscript.

## Supplemental Data

Supplemental figures and tables are available for this paper. Correspondence and requests for materials should be addressed to Laura Attardi.

